# Enabling inference for context-dependent models of mutation by bounding the propagation of dependency

**DOI:** 10.1101/2021.12.15.472813

**Authors:** Frederick A Matsen, Peter L Ralph

## Abstract

Although the rates at which positions in the genome mutate are known to depend not only on the nucleotide to be mutated, but also on neighboring nucleotides, it remains challenging to do phylogenetic inference using models of context-dependent mutation. In these models, the effects of one mutation may in principle propagate to faraway locations, making it difficult to compute exact likelihoods. This paper shows how to use bounds on the propagation of dependency to compute likelihoods of mutation of a given segment of genome by marginalizing over sufficiently long flanking sequence. This can be used for maximum likelihood or Bayesian inference. Protocols examining residuals and iterative model refinement are also discussed. Tools for efficiently working with these models are provided in an R package, that could be used in other applications. The method is used to examine context dependence of mutations since the common ancestor of humans and chimpanzee.

## Introduction

Early models of DNA sequence mutation assumed that each nucleotide was equally likely to mutate into any other nucleotide [Jukes and Cantor, 1969]. These were followed by others that incorporated differences in the mutation process, such as higher transition to transversion ratio [Hasegawa et al., 1985, Felsenstein and Churchill, 1996]. Although the rates of mutation of a nucleotide are known to depend on not only the nucleotide itself but also nearby sequence context [Seplyarskiy et al., 2021], it is difficult to take this context into account when performing inference.

This situation from biological sequence analysis is an example of a more commonly encountered class of models in statistics: a lattice of sites, with each site taking one of a finite collection of possible states, and whose stochastic, temporal evolution is Markov and governed by a set of local rules. For instance, at each site may sit a nucleotide whose mutation rate depends on physical properties determined by the nearby DNA sequence; a cell whose infection status depends on the state of its neighbors; or a particle whose spin is perturbed by external noise to a state whose distribution depends on the local energy configuration. Practical use of such models often requires inferring transition rules based on observations of the system at several time points, or of several states evolved along a tree from a single starting point. This paper concerns scalable methods for doing inference under such models using observations.

The situation is similar to the widely-studied problem of inference based on observation of a single instance of a Markov random field, and in many cases reduces to this if we only observe the system at one time point at stationarity. These are used, for instance, in spatial statistics [Besag, 1972, Gelfand et al., 2010] and image reconstruction [Geman and Geman, 1984, Besag, 1986]. The conditioning method we consider here is similar to the “coding” scheme introduced by Besag [1974], that conditions on a set of sites that makes the remaining observations independent; consistency of such methods has been shown by Comets [1992] and reviewed by Larribe and Fearnhead [2011]. A good review of recent statistical techniques is given by Friel [2012].

In the context of genomics, it is well-known that certain short nucleotide sequences are in many organisms much more, or less, abundant than expected by chance [Burge et al., 1992], due to the combined effects of context dependence of the nucleotide mutation process, selective constraints on the function of the sequence, and other processes such as biased gene conversion [Duret and Galtier, 2009, Arbeithuber et al., 2015]. Indeed, molecular studies have demonstrated that the spectrum of new mutations in humans is highly context dependent [Schaibley et al., 2013, Gao et al., 2019, Rodriguez-Galindo et al., 2020] and that multinucleotide substitutions are relatively common [Schrider et al., 2011, Terekhanova et al., 2013, Harris and Nielsen, 2013]. In humans, the molecular processes underlying mutations are best studied in contexts of intense somatic mutation: cancer and the hypervariable regions that generate immune system diversity [Cobey et al., 2015, Heredia-Genestar et al., 2020], which provide quite different cellular contexts but can share commonalities due to shared molecular machinery. In general, substitution rates depend on sequence context due to the interaction of a great many factors including: (a) likelihood of DNA damage or errors in cell division (e.g., UV damage [Goodman, 2002] or crossing over in meiosis [Arbeithuber et al., 2015]); (b) activity levels and structural properties of the enzymes responsible for repair (e.g., Y-family polymerases [Goodman and Woodgate, 2013, Sale et al., 2012]); and (c) any functional effects of an error (e.g., lethal mutations will not be seen in a population census). We are concerned with effects that are *homogeneous* across the sequence, so do not consider further the many constraints on protein-coding sequence [reviewed in Thorne, 2007]. Some of these effects – particularly, activity levels of different enzymes in the germline – will likely vary over evolutionary time, and so comparisons between widely separated species will identify time-averaged mutation rates. Studies have used a variety of exploratory techniques to identify sets of mutational patterns co-occurring in comparisons (a) between different cancer types [Alexandrov et al., 2013b, Shiraishi et al., 2015] and (b) between different human populations [Harris, 2015, Harris and Pritchard, 2016, Mathieson and Reich, 2017], to disentangle distinct signals presumably coming from the action of distinct sources of mutation.

However, phylogenetic and population genetic methods usually ignore such dependencies in the interest of computational efficiency, but some progress has been made. Often, the effects of a complex mutational spectrum are explored using summary statistics of larger blocks or other sensible but *ad hoc* methods. For instance, Arndt et al. [2003] studied dinucleotide transitions, and Yaari et al. [2013] displayed strong heterogeneity in the probability of synonymous mutations across 5-mers in B-cell immunoglobulin genes. Others have made progress using simplifying assumptions [Bérard and Guéguen, 2012], or by other approximations [Christensen et al., 2005, Saunders and Green, 2007]. Pedersen and Jensen [2000], later extended by Hwang and Green [2004], Hobolth [2008] and Baele et al. [2010a], used data augmentation and an MCMC algorithm to do Bayesian inference; Lunter and Hein [2004] used an approximate matrix decomposition; while Siepel and Haussler [2004] and later [Baele et al., 2010b] computed likelihoods by assuming a model that is Markov along the genome. However, all these methods are quite computationally intensive.

Another canonical example of context-dependent transition is the Ising model of statistical physics with time evolution given by Glauber dynamics [Glauber, 1963], in which a lattice of up/down spins are perturbed by thermal noise, relaxing into states dependent on the energy of the resulting configuration. Parameter estimation for Ising model without temporal dynamics is relatively well-understood [Pickard, 1982, Frigessi and Piccioni, 1990], but the problem of dynamical observations is less well-studied.

The general framework also fits certain cellular automata models, e.g., modeling wildfire [Clarke et al., 1994], the spread of HIV [Zorzenon dos Santos and Coutinho, 2001], or land use patterns [Wu, 2002]. Complex models may introduce long-range dependencies beyond the scope of this paper, but these methods may still prove useful in the modeling process.

This general class of models are known in the probability literature as interacting particle systems [Liggett, 2005] with neighborhood structure – continuous-time Markov chains on lattice-indexed collections of states whose transition probabilities are *local* in the sense that any instantaneous change only affects a small number of nearby sites, and the rates of such instantaneous changes depend only on the states in some bounded neighborhood of those sites to change. As finite-state Markov chains, transition probabilities are in principle simply expressable as a matrix exponential, but this is impractical because the size of the matrix is equal to the number of possible configurations.

In this paper, we propose a solution to this problem, showing that the conditional likelihood of local pattern count statistics can be well-approximated by marginalizing over a finite amount of surrounding context. We then use this to conjecture an approximation for the full likelihood. Motivated by the problem of inferring context-dependent mutation rates from diverged nucleotide sequences, we extend the inference framework to observations on trees. The tools are available as an R package, which is fairly efficient thanks to computational techniques using sparse matrices.

Before diving into a formal problem specification, we offer an intuitive explanation of how the method works. The difficulty of context-sensitive models is that, in principle, one may have a sequence of events with cascading consequences that involves a lot of the sequence. In the extreme case, one may have a sequence of mutations from, say, the left end of the sequence to the right end of the sequence, such that the probability of each mutation has been changed by that of the previous one.

However, the starting point of the work presented here is that for typical models such a sequence of events is improbable. For that reason, we can compute the transition probability for each segment of the sequence using a local context which, although bigger than the context which determines the local transition rates, is much smaller than the entire sequence. We call the building blocks of this computation *T-mers* (Figure 1) which parameterize transition probabilities for a small section of sequence (the *base* of the T-mer) based on additional sequence (the *overhang* of the T-mer). We can then bound the probability that the influence of a mutation outside the overhang of the T-mer will impact the sites at the base of the T-mer, which translates into an estimate of approximation error in our calculation of the likelihood.

**Figure 1:**
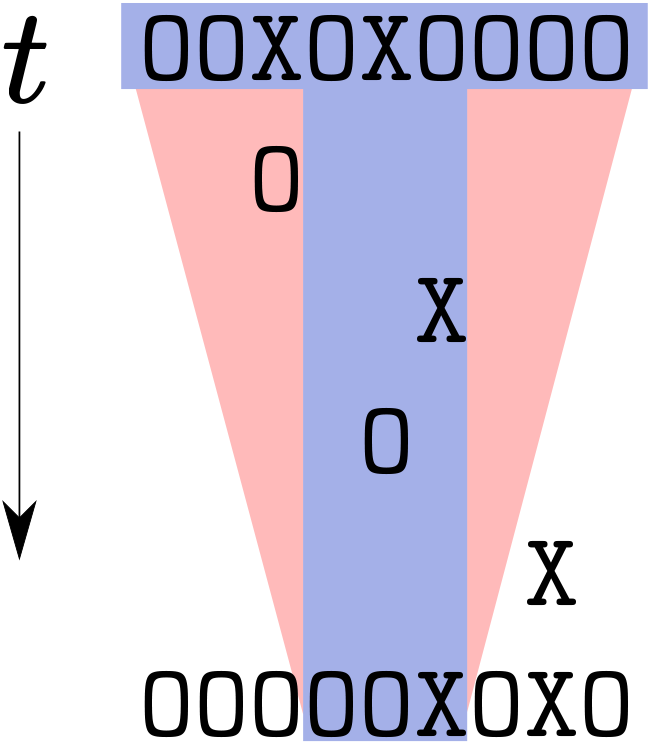
An overview of the method for a neighbor-dependent model, showing an initial sequence on top, a collection of mutations progressing through time, and a final sequence. The pink trapezoidal region represents the “range of influence”, i.e., possible locations and times at which a mutations is likely to affect the mutation outcome at the sites of interest (the three sites in the center of the final sequence). The blue T shape is called a *T-mer,* which is how our approximation to the process is parameterized. The central, smaller segment of the T is called the *base* of the T-mer, and the key building block computes the transition probability for this subsequence conditioned on the ancestral state at the additional *overhang* sites of the top of the T.

## 1 A general context-dependent model

Consider a 1-dimensional grid of sites, each of which can take one of a finite set of states, and that switch randomly between states according to a local set of rules. We then observe a finite collection of these sites at only a few times, Suppose that the dynamics are Markov: writing 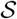 for the set of possible states and *X_i_*(*t*) for the state of site *i* at time *t*, we assume that {*X*(*t*)}_*t*≥0_ is a Markov process on sequences of *L* states 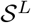 for which the probability a given site changes state in a small amount of time depends only on the sequence of states at nearby sites.

To formalize this notion of local dependence, first define a *pattern* to mean a contiguous sequence of states, and let |*u*| denote the length of a pattern *u*. Given a sequence 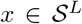, let 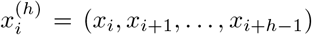 denote the pattern of *x* of length *h* beginning at location *i*. The dynamics of our stochastic process are determined by the set of allowed transitions and associated rates: given two patterns *u* and *v* of common length *h*, saying “pattern *u* changes to *v* at rate *μ*” means that

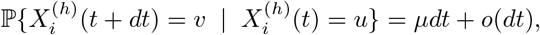

regardless of the location *i* along the sequence. If more than one pattern would match to cause the same change, their rates add. Such a rule is defined by its “transition triple” (*μ, u, v*), written as “*u* → *v* at rate *μ*”. A model of context-dependent mutation is defined by a set of all transition triples, which we denote 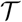.

Although our main focus is on nucleotides, it is informative to consider other, simpler, models as well. Here is one such example.

### Example 1.1

(TASEP). In the *Totally Asymmetric Simple Exclusion Process* describes a set of particles moving among some sites, with at most one particle per site. Each site is either “empty” or “occupied” (0 or 1, respectively), and each particle, independently at rate λ, checks whether the site to the right is empty, and if so, moves there. This process therefore has only one transition triple:

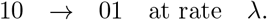

This only has one parameter. If we allow particles to exit from the right and new particles to enter from the left (e.g., by adding special symbols for the start and end of the sequence), then from observing only starting and ending configurations we cannot tell with certainty which particles have moved where, so it is not obvious how to estimate the speed. However, summing over possible particle movements would allow us to compute the likelihood of a given configuration change, and so estimate the speed by maximum likelihood given an initial sequence, a final sequence and an elapsed time.

To allow more compact specification of models, we also introduce a “potential”: suppose that there is a collection of patterns {*p_i_*} such that each occurrence of *p_i_* adds a quantity *e_i_* to the total “energy” of a sequence, and that the rate at which each possible change occurs is modulated by a function of the energy difference the change would produce. This is natural if transition triples describe *proposed* changes, and that the probability a proposed change actually occurs depends on how much it affects the energy.

Concretely, let 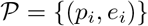 be a set of (pattern, energy difference) pairs, and for any sequence *x* define *E*(*x*) ∑_*i*_ *e_i_ n*(*x,p_i_*), where *n*(*x,p_i_*) is the number of times the pattern *p_i_* occurs in *x*. These will affect the rates through a nonnegative function *ϕ*: we declare that if the rate at which *x* changes to *y* as computed from transition triples is *μ*, then the actual rate is

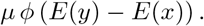

If *ϕ*(*e*) is not greater than 1, one way to think about this is that the transition triples give “proposed changes”, but these changes only take effect with probability *ϕ*(*E*(*y*) – *E*(*x*)). We have described this idea in terms natural for a physics model in which high energy configurations are disfavored, however it is also suitable for a population genetics model in which *ϕ* describes the probability of fixation of a mutant allele given a certain fitness change from the parent.

### Example 1.2

(Genomic GC content). A genome sequence can be written using A, C, G, and T; in the most general model of independent mutation across sites, each of the 12 possible transitions occurs at its own rate. Furthermore, in many species adjacent, methylated CG dinucleotides (“CpG sites”) have a much higher mutation rate to TG and CA than either single nucleotide change under the independent model. Embellishing the single-nucleotide model with this additional rate results in the model defined by

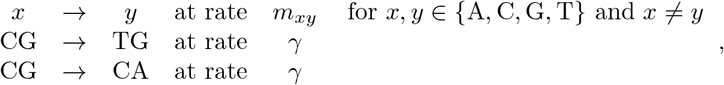

where *γ* is the additional CpG rate above the base mutation rate. Note that for a given pair of sequences, a change ACG → ATG could have occurred by a C → T mutation via the independent-sites model, or by a CG → TG mutation; in such cases the total instantaneous rate at which ACG changes to ATG is the sum of the rates (in this example, *γ* + *m*_CT_).

Counteracting this trend towards increased G/C is GC-biased gene conversion [Glémin et al., 2015], which acts effectively as a selective pressure against G or C bases. We model the evolution of a *single* sequence, not a population of sequences, imagining this sequence to be the consensus sequence of the population. A new mutation with an effect *s* on fitness that occurs in a population of effective size *N_e_* becomes ubiquitous, rather than dying out, with probability approximately (1 – exp(–2*s*))/(1 – exp(–2*sN_e_*)) [Kimura, 1962]. If the mutation occurs at rate *μ* per individual, the total rate it occurs at in the population is *μN_e_*. GC-biased gene conversion effectively means that sequences with more G/Cs are selected against. This is incorporated as follows: set the energy of patterns C and G to be *s*, so that *E*(*y*) – *E*(*x*) is the net change in number of G’s and C’s multiplied by *s*. Also, define *ϕ*(*e*) to be the expected rate of fixations of “selected” mutations with advantage *e* relative to the per-individual, per-site mutation rate:

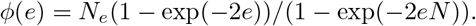

With these definitions, if a given change that occurs with rate *μ* would change a sequence *x* to *y*, then *μϕ*(*E*(*y*) – *E*(*x*)) is the rate at which the mutation appears and successfully takes over in the population.

We close with a final example from statistical physics, to be used later:

#### Example 1.3

(Gibbs sampling of the Ising model). In the Ising model, each site is labeled as either “up” or “down” (+1 or −1 respectively), imagined as a string of n magnetic dipoles, and the energy associated with a given state *x* is 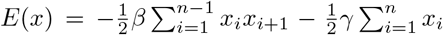. Here *β* represents inverse temperature, and *γ* represents the strength of the magnetic field (here scaled by temperature). The associated stationary distribution on configurations is proportional to exp(−*E*(*x*)). The following process, also known as “Glauber dynamics”, preserves the stationary distribution: each site, independently at rate λ, forgets its spin, and reconfigures to a state chosen with probability proportional to the stationary probability of the resulting configuration: Since exp(−*E*(*x*))/(exp(−*E*(*x*)) + exp(−*E*(*y*))) = 1/(1 + exp(−(*E*(*x*) ‒ *E*(*y*))), this is:

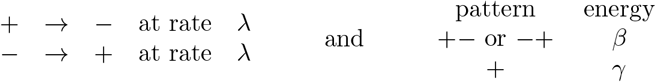

and

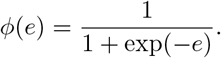

### The generator matrix

We now describe concretely how the set of transition rates on patterns determines the transition rate matrix for complete sequences, that is, the 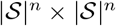 matrix *G*(*n*) whose (*x, y*)^th^ entry gives the instantaneous rate with which the process in state 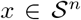 jumps to state 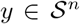. Recall that 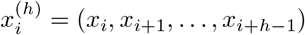 denotes the subsequence of length *h* beginning at location *i*. For each 1 ≤ *i* ≤ *n*, pattern length *h* > 0, and patterns 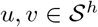 define the relation

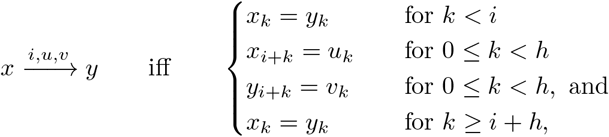

i.e., if *x* and *y* match except at positions *i, i* + 1,…, *i* + *h* – 1, and for those positions *x* matches with *u* while *y* matches with *v*.

As before, let 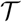 be the set of transition triples for the model. As noted above, there may be more than one way to mutate a sequence *x* to get *y*: for each position *i*, let *J*(*i, x, y*) be those transitions in 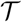 that can be applied at *i* to change *x* into *y*, i.e.,

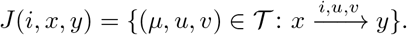

The rate *G*(*n*)_*x, y*_ is then the sum of all transition rates that can take *x* to *y*, multiplied by the fitness term, namely

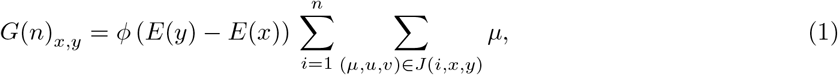

and if there are no triples (*μ, u, v*) with 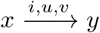 for some *i*, then *G*(*n*)_*x,y*_ = 0. To make this the generator matrix for a Markov process, we also set each *G*(*n*)_*x, x*_ so that rows sum to zero. Indeed, this matrix is very sparse because for most pairs (*x, y*) there are no transition triples that can change *x* into *y*.

In principle, this gives us the transition probabilities for the process by a matrix exponential:

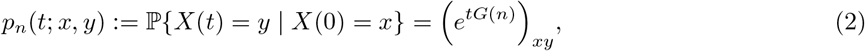

which would then provide a route to parameter estimation. In practice, the size of these matrices makes direct application obviously impractical for anything but very small *n*. This paper proposes an alternate strategy.

## 2 Simulation

First, it will be useful for later proofs to have an explicit construction of the process, also used for simulation, using a variant of the “jump chain representation”, also known as the “Gillespie algorithm” or “uniformization” [Hobolth and Stone, 2009]. Briefly, this works by first sampling a homogeneous Poisson process of times and locations of possible changes, at an appropriate rate *μ*\* per site, and then resolving each possible change in temporal order. (Note that these “possible changes” are different than the “proposed changes” described above in terms of the energy model.) The rate *μ*\* should be the maximum “local” rate at which transitions occur, across sites and transition outcome states:

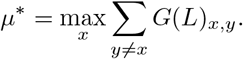

Note that we do not actually need to construct *G*(*L*) to find the maximum rate. Now sample possible changes: in a sequence of length *L*, over a time period *t*, first draw the number of possible changes, *N*, from a Poisson distribution with mean *μ*tL*, and then choose the times *s_k_* and locations *i_k_* at which possible changes occur uniformly over [0,*t*] and {1, 2,…, *L*} respectively. Order the possible change times, so that

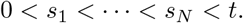

(Note that this is equivalent to saying that possible changes occur at rate *μ*\* independently at each of the *L* sites.) Now, suppose we have determined the state just before time *s_k_* to be *X*(*s*_*k*–1_) = *x*. Then, the new state *X*(*s_k_*) is chosen by applying a mutation at position *i_k_* with probability proportional to the corresponding rate, i.e., out of the possible transitions 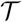, transition triple (*μ*(*j*),*u*(*j*),*v*(*j*)) is chosen with probability

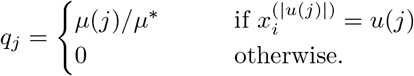

The remaining probability 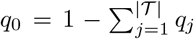 gives the probability that the state remains the same. (By construction of *μ*\*, *q*_0_ ≥ 0.) If triple (*μ*(*j*),*u*(*j*),*v*(*j*)) is chosen, then the substring of *x* matching *u*(*j*) that begins at position *i_k_* is replaced with *v*(*j*) to produce *X*(*s_k_*).

For instance, to simulate the TASEP, one has only to let *μ*\* = λ, and at each possible change check if at site *i_k_* there is a 1 followed by a 0, and if so, switch them.

## 3 Inference

The problem at hand is to infer the parameters of the model based on the final state of the model after it has evolved from some known initial state. How can we extract the information? These are Markov processes, on the state space 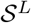, so the full likelihood function is given by (2), but doing anything with the 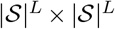 matrix *G_L_* is clearly infeasible even for moderately sized *L*. We can, however, compute (2) with smaller *G_n_*, so the first thing that one might think to do is to break the sequence up into many blocks of length m, and treat these as independent. Then, defining *p_m_*(*t*; *x, y*) to be the probability that a string x of length m evolves to string y over time t, we would obtain the approximate likelihood function

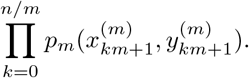

However, this ignores dependencies between neighboring blocks, so it is not clear how good this approximation might be.

The key insight we use is that by including neighboring sequence context in the *initial* sequence only, we can compute *something* correctly, and that will be sufficient to compute the full likelihood, at least arbitrarily. To do this, let *p_n,ℓ,r_*(*x, y*) for ℓ + *r* < *n* denote the probability that an initial subsequence *x* of length *n*, after time *t*, is found to match a smaller subsequence *y* of length *n* ‒ ℓ – *r*, offset by ℓ sites (see Figure 2). This central, smaller subsequence is called the “base” of the T-mer. We refer to such patterns as *T-mers*, as depicted in Figure 2. We emphasize that the overhang of the T-mers is typically bigger than the context-sensitivity width of the underlying model (Figure 1).

**Figure 2:**
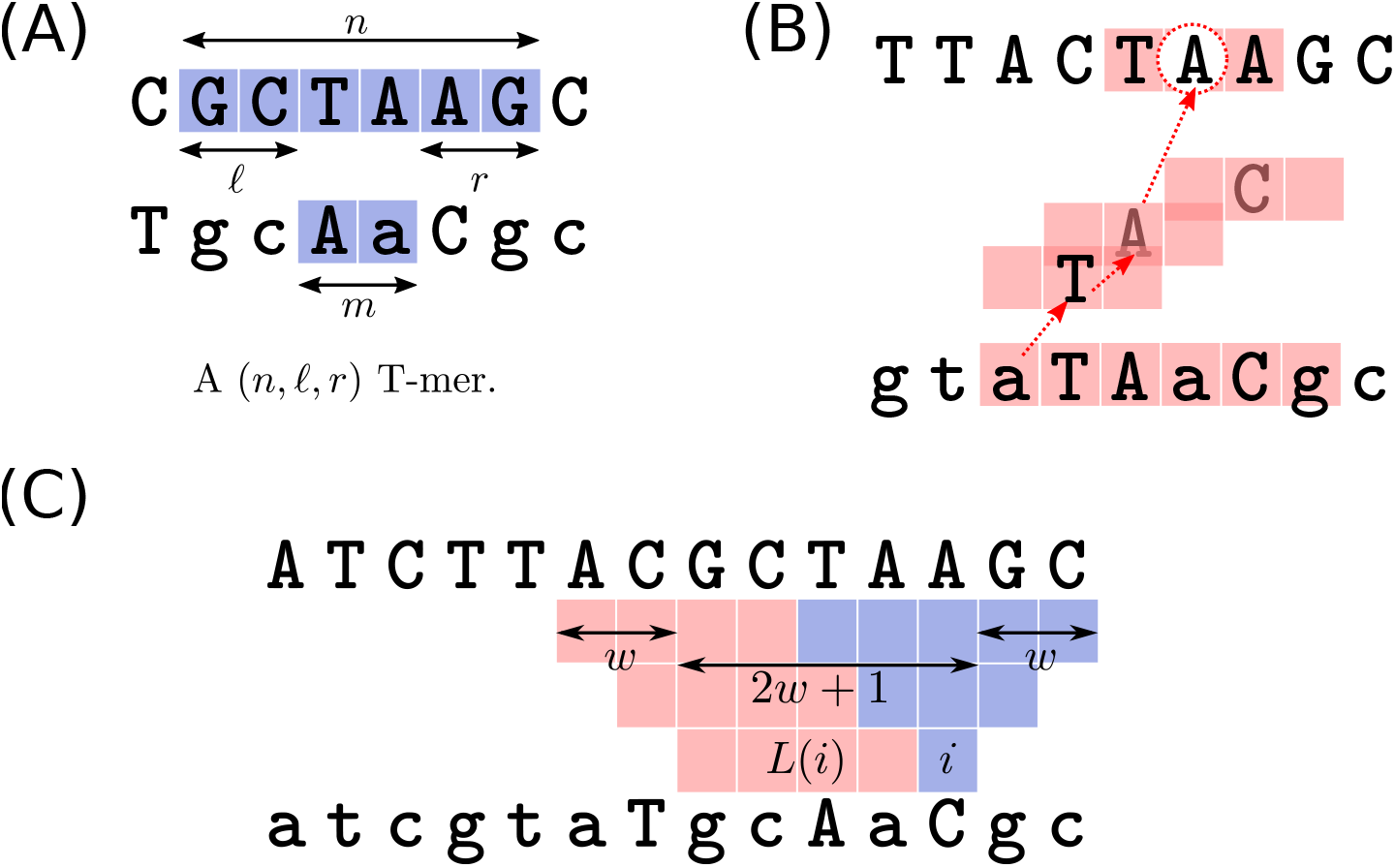
**(A)** A (6, 2, 2) T-mer. Nonmutated sites are shown in lower case letters. The “base” of the T-mer is the lower central string of length *m* = *n* – ℓ – *r*. **(B)** The three mutations have extended the influence of the circled A in the initial sequence (top) to the six positions colored red in the final sequence. The rate of each mutation depends on its’ neighboring sites, i.e., with a window of width *w* =1. Red dotted arrows show the propagation of dependency between the circled A and the third site in the final sequence (bottom). **(C)** Diagram of heuristic argument for full likelihood approximation: site *i* depends on the blue squares in *N_i_*; the sites to the left of this in *L*(*i*) are those other sites in *X*(*t*) whose dependency neighborhoods overlap *N_i_*. Red squares give the dependency neighborhood of *L*(*i*), so we condition on the row of red and blue sites at the top. On the bottom, we have 2*w* + 1 sites; at the top we have 4*w* + 1; here *w* = 2.

These *p_n,ℓ,r_*(*x, y*) probabilities can be found by marginalizing *p_n_*, summing over the possible initial and final substrings of *x* (denoted *a* and *b*):

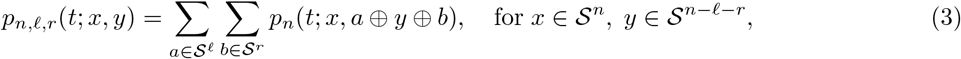

where *a* ⊕ *y* ⊕ *b* is the sequence composed of concatenating *a, y*, and *b* together in that order. We expect *p_n,ℓ,r_*(*x, y*) to be asymptotically correct when ℓ and *r* are large (and *t* is fixed). In Appendix B, we show that this is so, namely that the probability a given subsequence of length *n* matching *x* is later observed to have its center matching *y* is close to *p_n,ℓ,r_*(*x, y*), regardless of wider context. Formally, for each *i*,

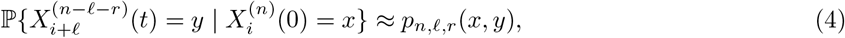

with the approximation getting better for longer ℓ and *r*.

Equation (3) could then be used to compute a composite likelihood, which we denote *A_n,ℓ,r_*(*x, y*). This is the product of the probabilities of transition for all (*n, ℓ, r*) T-mers between *x* to *y*, i.e., with *m* = *n* – ℓ – *r*,

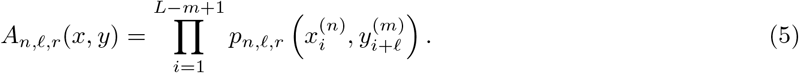

To keep notation simple we’ve ignored the boundaries of the sequence; to deal properly with these, on the right-hand side of the equation ℓ should be replaced with min(ℓ, max(*i* – 1,0)), and *r* should be replaced with min(*r*, max(*n* – *i* – ℓ – *m* + 1,0)).

### 3.1 Full likelihood

What we actually want is the *full likelihood* 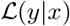, the probability that a given sequence *x* evolves into another, *y*, over a given period of time. This full likelihood can be computed as the product of the probability of each site in *y* given *x* and all sites in *y* further to the left. Defining *x*_<*i*_ = (*x*_1_,…, *x*_*i*–1_) to be the subsequence to the left of *i*, this is:

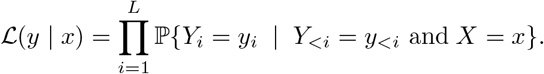

Each term in this product is the probability of *Y_i_*, given all sites in *X* and all sites in *Y* to the left of *i*. It seems reasonable that, similar to equation (4) above, the process at position i should be approximately independent of distant positions, given the neighboring positions.

Here is a heuristic argument, depicted in Figure 2C. Suppose that ℓ is long enough that approximation (4) holds with ℓ = *r*. Then we know that site i approximately depends only on mutations within distance ℓ. Furthermore, the sites up to distance 2ℓ to the left of *i* also depend on these mutations. We can now use (4) to restrict the dependency on *x*. Recall that the last base of 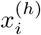 is *x*_*i*+*h*–1_. So, 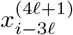 runs from *i* – 3ℓ to *i* + ℓ, inclusive, while 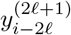 runs from *i* – 2ℓ to *i*, inclusive. This suggests that the following approximation should be good:

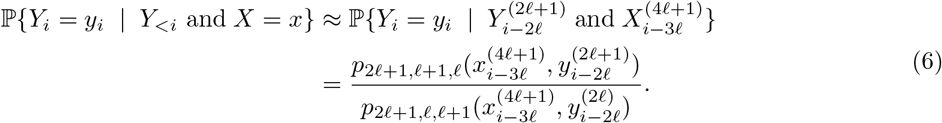

Since the full likelihood is a product across sites, we can then simply express an approximate full likelihood as a ratio of two composite likelihoods:

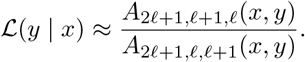

See the next section for more discussion of the approximations.

### 3.2 Choice of overhang

The approximations in (4) and (6) hold if the overhang, ℓ, is long enough. We present intuition about these formulas here, which provides a way to pick the overhang lengths ℓ and *r*; a precise statement is given in Appendix B. Suppose that each instantaneous change depends on positions at most *w* sites away, and refer to *w* as the “window width”. Then, the mutation process at any particular site depends on the initial sequence at that site and within distance *w* on either side… *and* the outcomes of any mutations that have occurred within that window (see Figure 2).

Therefore, the sequence at a particular site may directly affect the process at the neighboring *w* sites on either side. If another change happens, say, at the end of this window, then the indirect effects of this change may extend out to 2*w* sites. In this way, the influence of the sequence at each site extends outwards, as shown in Figure 2 – so if we take the overhangs long enough that this dependency is unlikely to extend to the base of the T-mer, the approximation will be good. As we show in Section B, if the overhangs are *t* = *r* = *kw*, then the error in approximation (4) is bounded by the probability that there are more than *k* changes in a given window of length *w*. Concretely, sites further away than *kw* only matter if there is a chain of at least *k* intervening mutations – which is unlikely if *k* is much larger than the expected number.

For instance, in the CpG model the window width is *w* = 1 because the process only depends on neighboring sites; and if the sequence has evolved for time *t* then we expect no more than a Poisson(λ*t*) number of changes at each site. If λ*t* = 0.75 (1 – *e*^-3/4^ = 52% of the sequence will have changed!), then taking *t* = *r* = 3 allows us to compute T-mer probabilities to within an error of 0.007, since this is the probability that a Poisson with mean 0.75 will be larger than 3. If λ*t* = 0.25 we would do even better with *t* = *r* = 2.

Appendix B makes this argument more precise. The argument for the full likelihood is more difficult, so at present our proposal above remains a conjecture. To see one difficulty, consider the TASEP model, with *x*_1_ = *y_n_* = 1, and all remaining positions equal to 0. The full probability of this pair of sequences is very small, while the conditional probability of *Y_n_* = 1 given *X* = 10 ⋯ 0 and *Y*_<*n*_ = 0 ⋯ 0 is large since the 1 that initially began in the first position had to go somewhere. However, if we do not include the first position in the conditioning, this conditional probability is very different. Fortunately, such situations are rare, and the TASEP model provides the extreme case of dependency propagation. In practice, we recommend computing the likelihood with equation (6) using successively larger values of t to check for convergence.

## 4 Computation

For inference, we need to compute the 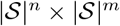 matrix *F* whose (*a, b*)^th^ entry is *p_n,ℓ,r_*(*x, y*), for various *t* (and *m* = *n* – ℓ – *r*). This matrix is a projection of the 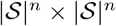 matrix whose (*a, c*)^th^ entry is

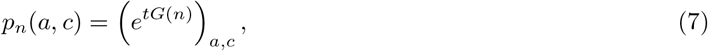

where *G*(*n*) is the sparse 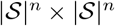 matrix defined in (1). The “projection” we need just marginalizes over long (i.e., length *n*) patterns *c* that match the shorter (i.e., length *m*) pattern *b* as in (3): if we define the *n* × *m* matrix *U* so that *U_cb_* = 1 if 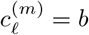 and *U_cb_* = 0 otherwise, then

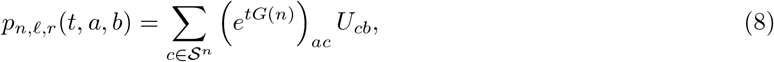

so in fact we don’t need the entire matrix *e*^*tG*(*n*)^, just the product of this matrix with each of the columns of *U*. Modern techniques in sparse matrix computation (e.g., Krylov methods) provide efficient ways to do this.

Using the R package expm [Goulet et al., 2017], this makes computation quite feasible: with four possible states and ℓ = 2 and *m* = 1, so that *G*(*n*) is a 1024 × 1024 matrix, computing *e*^*tG*(5)^ takes 13 seconds, while computing *e*^*tG*(5)^*U* takes only 0.3 seconds. Increasing to ℓ = 4 and *m* = 1 or ℓ = 3 and *m* = 2 is still feasible, taking *e*^*tG*(8)^*U* in 42 and 47 seconds, respectively (and much longer for the whole matrix *e*^*tG*(8)^).

### Sparsity and updating

*G* We can also take advantage of the sparsity of *G* to perform efficient computation of the likelihood under many sets of parameters. Let’s assume that the T-mers only allow single-position changes. First note that in this case *G*(*n*) has 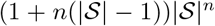 nonzero entries, since each of the 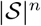 sequences can change in *n* places. This will determine how the computation scales with *n* and 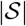, once we precompute a number of things. Let *g* = (*g*_1_,…, *g_d_*) be the nonzero entries of *G*(*n*), in some fixed ordering; sparse matrix representations of *G*(*n*) store only *g* along with information about the rows and columns these are found in. Each *g_i_* is a linear combination of mutation rates *μ_j_*, multiplied by a function (*ϕ*) of a linear combination of energy coefficients ej, say *g_i_* ∑_*j*_ *A_ij_μjϕ*(∑_*k*_ *B_ik_e_k_*), so by precomputing the matrices *A* and *B* we can update *G*(*n*) with new parameter values using only two matrix multiplications and evaluation of *ϕ*() over a vector. This remains efficient to perform in interpreted languages such as R, since each step is carried out by lower-level compiled code (e.g., optimized linear algebra libraries).

## 5 Phylogenetic inference

In phylogenetic applications, rather than “before” and “after” observations, we get two (or more) observations evolved from a common root. In the simplest case of two tips we have two processes *X* and *Y*, with identical starting states *X*(0) = *Y*(0), observed only at times *t_X_* and *t_Y_* respectively. At first, one might think to compute probabilities with a “long” sequence at the root and “short” sequences at every tip. However, this does not work out, since the root is unobserved and short sequences at every tip give us no information about what the root sequence in the overhangs might be.

Instead, we will compute the probability that we see (long) pattern *x* in *X*(*t_X_*) juxtaposed with (short) pattern *y* in *Y*(*t_Y_*), summing over possible (long) patterns at the root. Concretely: pick a random location *I* in the sequence and let 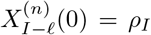 be the (long) pattern seen there at the root. Write *π*(*z*) for the frequency of a given pattern *z* seen there, i.e., 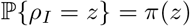. The probability that we see *x* and *y* at the random location *I* is

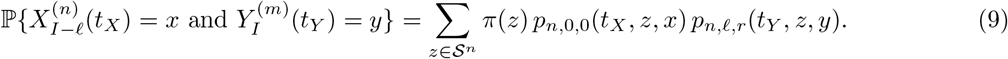

Denote this quantity *p_n,ℓ,r_*(*x, y*). This can be computed without much more effort than the simpler case above, given the frequencies at the root, as described in greater generality below. These calculations can be done using the same sparse matrix methods as above. For instance, to compute

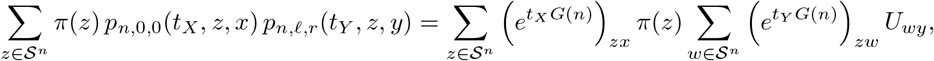

we can first compute 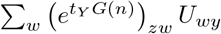 as above, multiply rows by *π*(*z*), and then matrix multiply by the transpose of *e*^*t_x_G*(*n*)^.

We might approximate the full likelihood as follows, after choosing *n, ℓ*, and *r* appropriately (see above). For the sake of notational simplicity, assume that we have a few positions of *X* observed before the start and after then end of the sequence (so that, e.g., we can refer to *x*_*n*+1_,…, *x_n+r_*).

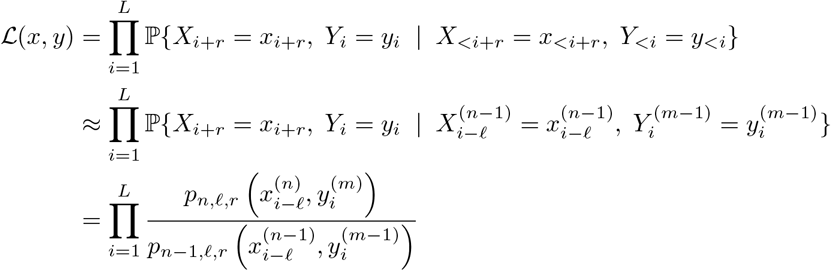

We expect this to be a good approximation to the true likelihood when the same conditions above are met, taking t to the be the largest distance between two taxa in the tree.

### 5.1 Phylogenetic pruning

The general case is a modification of Felsenstein’s “pruning” algorithm [Felsenstein, 1981].

To compute the likelihood on a more complicated tree, consider the following, with the three-taxon tree of Figure 3 as an example, where we are counting occurrences of (long) *n*-tuples in taxon *X*, and (shorter) *m*-tuples in taxa *Y* and *Z*. The likelihood in this example is found by summing over the root state (the summation over *u*) and the interior node (*v*):

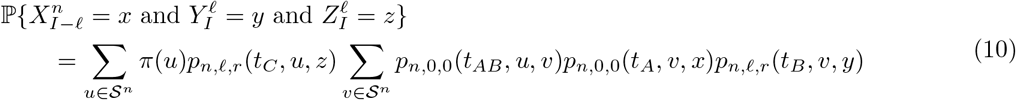

**Figure 3:**
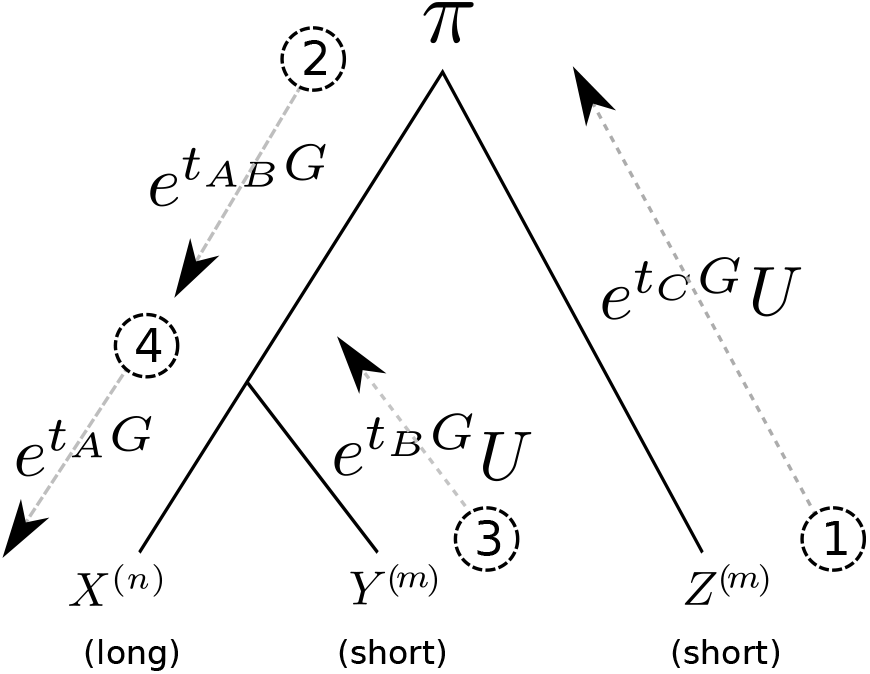
A depiction of the four steps in computing the likelihood of finding particular “short” subsequences at *Y* and *Z* given the “long” subsequence at *X*: see text for details.

The steps for computing this are then:

1. Compute *M*_1_ (*u, z*) = *p_m,ℓ,r_*(*t_C_, u,z*)(*a* 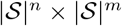 matrix)
2. Compute 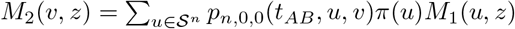 (still a 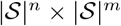 matrix)
3. Compute *M*_3_(*v, y*) = *p_m,ℓ,r_*(*t_B_, v, y*) (a 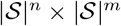 matrix)
4. Compute 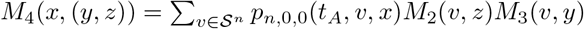 (stored as a 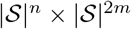 matrix).

Then *M*_4_ gives all the likelihoods. This is depicted in the figure, writing (*e*^*tG*^)_*ab*_ for the matrix *p*_*n*,0,0_(*t, a, b*), and *U* for the projection matrix that gives *p_m,ℓ,r_*(*t, a, b*) = (*e^tG^U*)_*ab*_.

In total there are 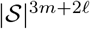 possible data combinations. However, we can reduce dimensionality by only computing probabilities for certain (more common) patterns: for instance, one could compute the likelihood for situations where *y* and *z* agree (but may differ from *x* by replacing step (4) with:

4′. Compute 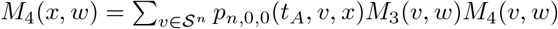 (stored as a 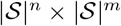 matrix).

On a three-taxon tree it is not clear that this is desirable, but on larger trees it seems likely that we’d want to restrict to combinations with most sites agreeing across the tree. (Note, however, that one should not first look at the data, observe which combinations (*y, z*) were most common, and then restrict to only those!)

The algorithm for a general tree is written out in Appendix C.

### 5.2 Model fit and model selection

Fitting a model to nucleotide data will likely require iterative modification of the model. The difference between observed and expected T-mer counts provide a good assessment of how and whether a model can be improved. These *residual* counts are usually most usefully computed for T-mers shorter than those used to fit the model, because the smaller total number of possible T-mers makes it easier to identify the shared patterns. For each T-mer (*x, y*), write *O_x,y_* for the number of this T-mer observed in the data, and *E_x,y_* for the number expected under the model. (The expected number is the number of occurrences of x in the initial sequence multiplied by the probability *p_n,ℓ,r_*(*t*;*x,y*), where *n* is the length of *x*, and *ℓ, r* are the appropriate overhangs.) If the T-mers were nonoverlapping, this would form a standard contingency table, and standard practice would be to divide the residuals by the square root of the expected counts and compare these to a standard Normal distribution. However, there are are n possible such tables – so, we compute normalized residuals as

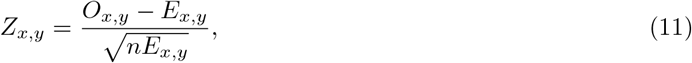

which is equivalent to averaging the *z*-scores of the *n* possible tables of nonoverlapping counts. These can then be compared to a standard Normal distribution.

## 6 Software

The methods described here are implemented in R and freely available at https://github.com/petrelharp/context. The core implementation of T-mer counting and likelihood evaluation is provided in an R package, `contextual`, and is supplemented by (documented) code to simulate from context-dependent models and perform maximum likelihood or Bayesian inference.

## 7 Results

### 7.1 Stepwise model selection

We benchmarked the performance of stepwise model selection: (a) fit a simple model, (b) choose new motifs to add based on statistically significant residuals, (c) repeat until no residuals are statistically significant. To test this method, we simulated a single sequence of 10^6^ nucleotides for one unit of time under the model given in Table 1, obtained by embellishing the basic CpG hypermutability model with the longer mutational motif identified by Harris [2015]. The resulting sequences differed at 28.5% of the sites.

To begin, we fit an unconstrained non-context-dependent model of single-nucleotide substitution using a constrained quasi-Newton method (`optim(.., method=‘L-BFGS-B’)` in R). Recall that for inference, we next need to pick a T-mer shape with which to compute likelihoods, and the shape of this T-mer is only determined by the shape of dependencies in the generative model in that it determines the minimum base and overhang lengths. We use (5,1,1) T-mers: three sites at the base with an overhang of one on each side. Since this initial model has no context-dependence, we could do the first step of model fitting using (1,0,0) T-mers (i.e., single positions), but for simplicity we use the same T-mer shape for all steps.

After fitting the model, we can look at residuals using any T-mer shape we like. At this first step, we computed the (2,1,0) and (2,0,1) T-mer residuals, shown in Table 2. The largest residuals are an excess of C → T mutations in CG dinucleotides, and the reverse-complement pattern (*z* scores of 88). Complementing this are deficits of C → T mutations in other contexts, since the C → T mutation rate has been fit to be an average across contexts. This clearly calls for addition of the CG → TG mutation motif (and reverse complement), which was used in simulating the data.

After adding these patterns into the model and re-fitting, we searched for evidence of additional processes happening in the data. To do so, we calculated length 2 residuals and (3,1,1) T-mer residuals. Although some of the length 2 residuals had large *z*-scores, none were as large as those seen in (3,1,1) T-mer residuals, which are shown in Table 3. These show an excess of TCC → TTC and AGG → AAG changes, which were indeed the motifs used when simulating the data. After adding these in, no further (3,1,1) T-mer residuals achieved statistical significance after computing *p*-values with the Gaussian CDF and applying a Bonferroni correction. The final parameter values are shown in Table 1; parameter estimates differ from the truth by less than 3% (mostly within 1%).

**Table 1:** True parameter values, and MLE estimates obtained from iterative fitting as described in the text to a 10^6^ bp sequence evolved to have 28.5% sequence divergence with the above mutational motifs. The transitions CG → TG and CG → CA were constrained during fitting to be equal (DNA strand symmetry), as were the transitions TCC → TTC and AGG → AAG.

| x | → | y | at rate | truth | MLE |
| --- | --- | --- | --- | --- | --- |
| A | → | T | at rate | 0.10 | 0.0996 |
| T | → | A | at rate | 0.10 | 0.0982 |
| C | → | G | at rate | 0.15 | 0.1505 |
| G | → | C | at rate | 0.15 | 0.1497 |
| A | → | C | at rate | 0.08 | 0.0784 |
| T | → | G | at rate | 0.08 | 0.0792 |
| A | → | G | at rate | 0.08 | 0.0798 |
| T | → | C | at rate | 0.08 | 0.0795 |
| C | → | A | at rate | 0.12 | 0.1202 |
| G | → | T | at rate | 0.12 | 0.1196 |
| C | → | T | at rate | 0.12 | 0.1198 |
| G | → | A | at rate | 0.12 | 0.1193 |
| CG | → | TG | at rate | 0.40 | 0.3978 |
| CG | → | CA | at rate | 0.40 | 0.3978 |
| TCC | → | TTC | at rate | 0.10 | 0.1033 |
| AGG | → | AAG | at rate | 0.10 | 0.1033 |

### 7.2 Bayesian inference

The usage of fast sparse matrix methods described above make likelihood computation fast enough to use a Metropolis-Hastings-based Markov chain Monte Carlo method to sample from the posterior distribution on parameters. To test this, we evolved 10^6^ sites from the Ising model (described above) with λ = 0.5, *β* = 1.0, and *γ* = 0.5 for 1 unit of time, resulting in around 460,000 possible mutations and differences at around 186,000 sites between initial and final sequence. We then placed an Exponential prior with mean 1 on both transitions +–→ – and → + (without constraining these to be equal) and half-Gaussian priors with mean 0 and scale 3 on both *β* and *γ*, counted (5, 3,1) T-mers, and used the `mcmc` package [Geyer and Johnson, 2017] to run a random walk sampler for 10,000 steps (judged sufficient by examination of diagnostic plots). We repeated this procedure separately on 61 independently simulated sequences, and show the resulting credible intervals in Figure 4.

**Figure 4:**
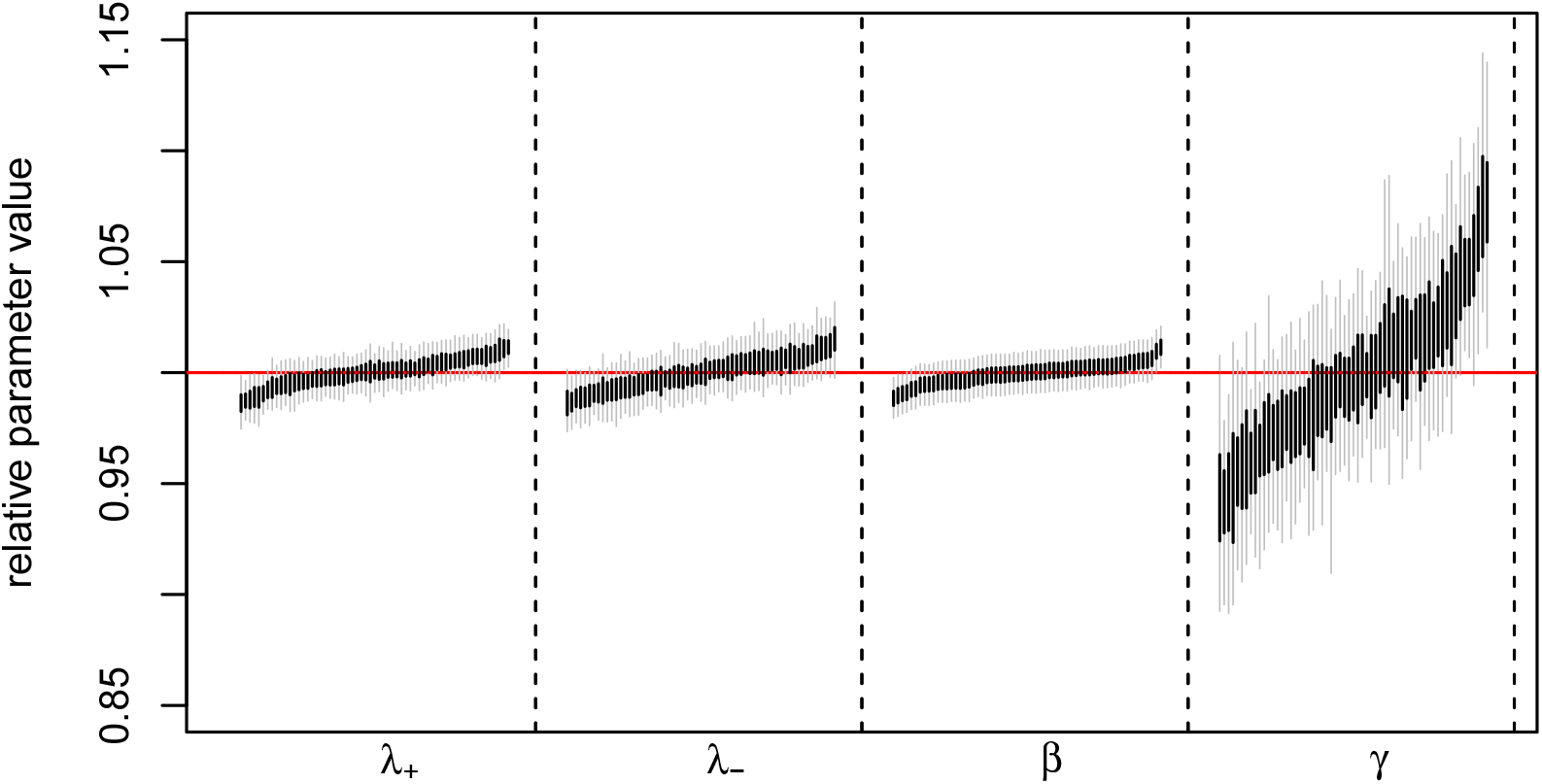
Credible intervals for the four parameters of the dynamic Ising model, calculated using (6, 2, 2) T-mers, from 61 MCMC runs on independently simulated sequences of 10^6^ bases differing at around 18.6% of the sites. Dark bars show the 50% credible intervals, and lighter grey bars the 95% credible intervals. Values shown are relative parameters, obtained by dividing by the true value.

Posterior medians were within half a percent of the truth for J → – and → + (corresponding to λ) and the temperature (*β*), but showed around 2% error for the magnetic field *γ*. As seen in Table 4, credible intervals were somewhat too narrow for shorter T-mers, but (6, 2, 2) T-mers were long enough with sufficient overhang to produce well-calibrated posteriors.

### 7.3 Hominid mutation spectrum

We also applied the method to human–chimpanzee divergence data, restricting the comparison to putative regulatory sequences, avoiding the additional constraints of coding sequences. To do this, we extracted regions in the human (hg38)-chimpanzee (panTro4) alignment according to four types of regulatory feature in the Ensembl Regulatory Build [release 81 Zerbino et al., 2015]: “CTCF binding site”, “enhancer”, “open chromatin region”, and “promoter flanking region”. These were furthermore divided into regions that either did or did not overlap a transcription start or end site of a known gene from the hg38 Annotations Database [Kent et al., 2002] and filtered for length: aligned regions below either 3000bp (promoter flanking regions) or 1000bp (all other types). We then counted T-mers of appropriate length in each of the eight resulting sets of aligned regions, omitting any T-mers that included gaps or repeat-masked positions, and fit models separately to each of the eight data sets.

**Table 2:**
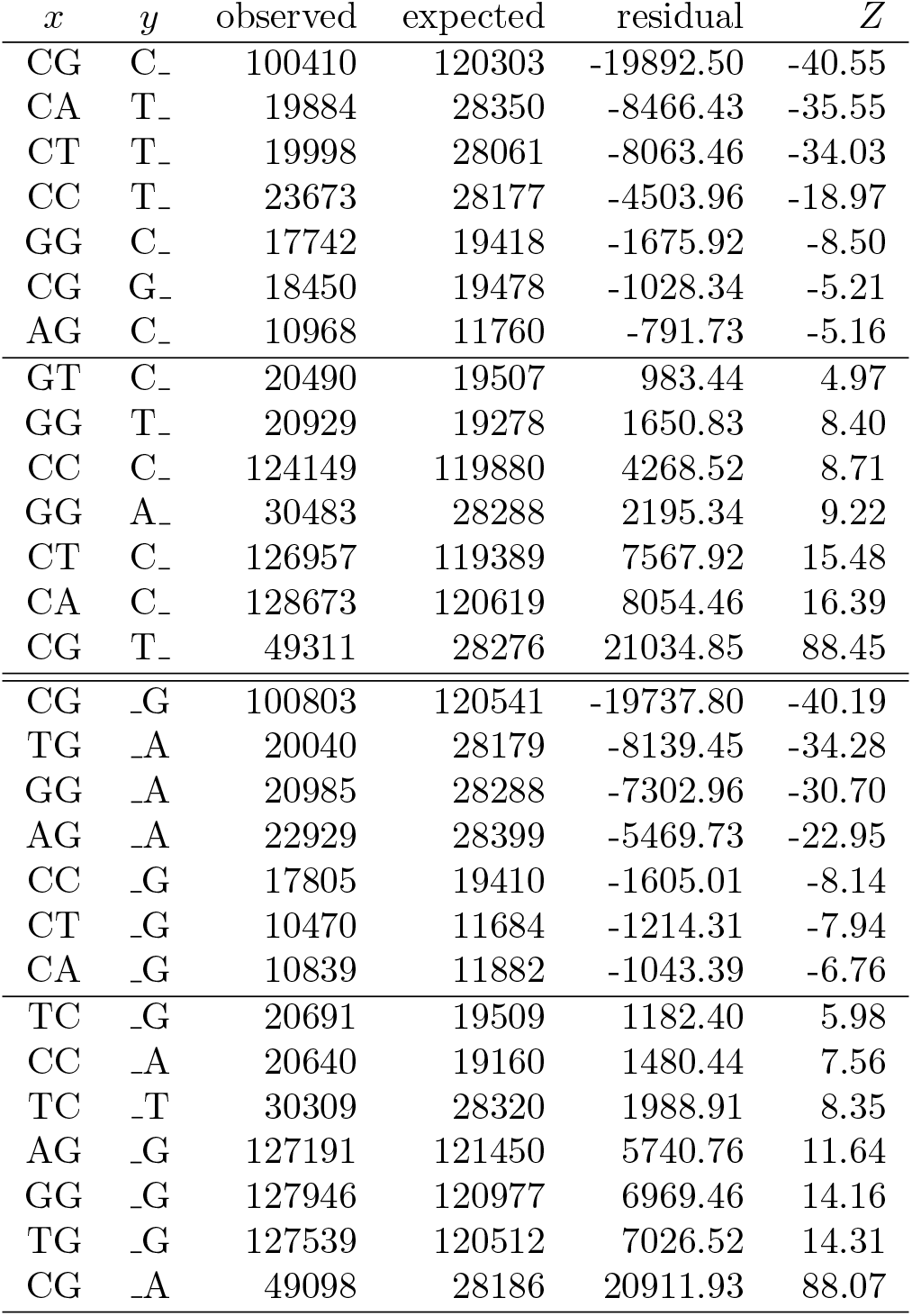
Largest residuals of shape (2,1,0) and (2,0,1) after fitting a model with only “basic” (singlenucleotide) mutations to sequence simulated under the model of Table 1, that also includes CpG hyper-mutability as well as TCC → TTC and AGG → AAG mutations. Note that in the table, the underscore means “any nucleotide”.

**Table 3:**
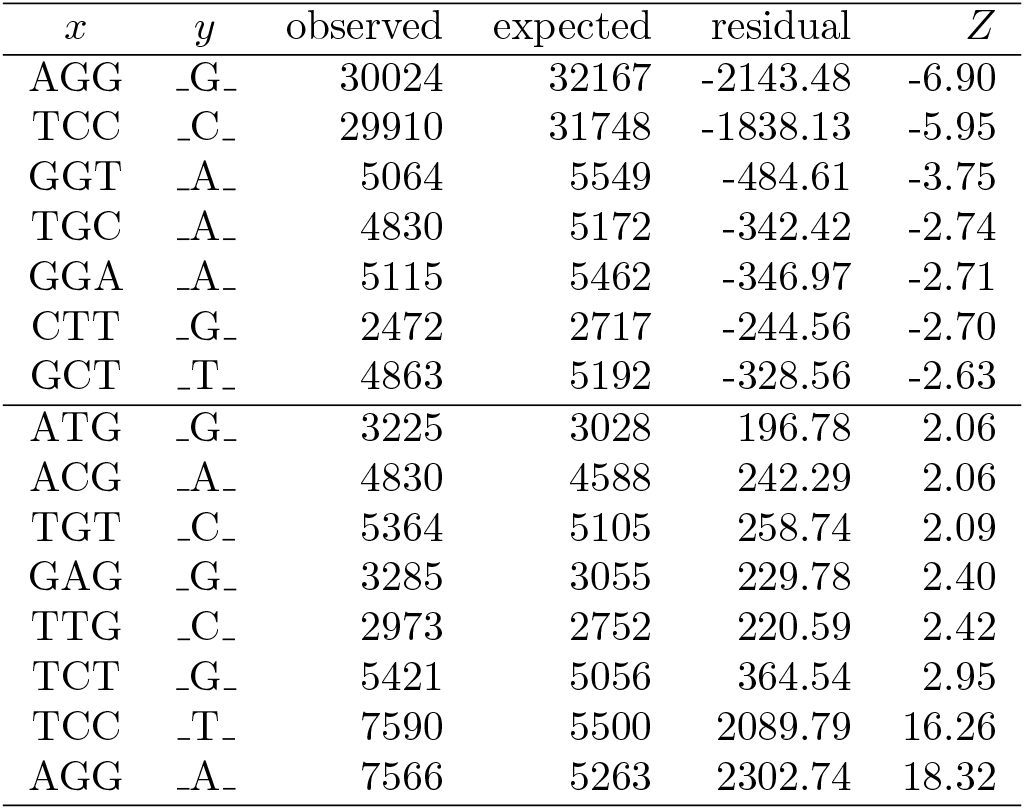
Largest residuals of shape (3,1,1) after fitting a model with “basic” (single-nucleotide) and CpG mutations to sequence simulated under the model of Table 1, that also includes TCC → TTC and AGG → AAG mutations. Note that in the table, the underscore means “any nucleotide”.

| $x$ | $y$ | observed | expected | residual | $Z$ |
| --- | --- | --- | --- | --- | --- |
| AGG | _G_ | 30024 | 32167 | -2143.48 | -6.90 |
| TCC | _C_ | 29910 | 31748 | -1838.13 | -5.95 |
| GGT | _A_ | 5064 | 5549 | -484.61 | -3.75 |
| TGC | _A_ | 4830 | 5172 | -342.42 | -2.74 |
| GGA | _A_ | 5115 | 5462 | -346.97 | -2.71 |
| CTT | _G_ | 2472 | 2717 | -244.56 | -2.70 |
| GCT | _T_ | 4863 | 5192 | -328.56 | -2.63 |
| ATG | _G_ | 3225 | 3028 | 196.78 | 2.06 |
| ACG | _A_ | 4830 | 4588 | 242.29 | 2.06 |
| TGT | _C_ | 5364 | 5105 | 258.74 | 2.09 |
| GAG | _G_ | 3285 | 3055 | 229.78 | 2.40 |
| TTG | _C_ | 2973 | 2752 | 220.59 | 2.42 |
| TCT | _G_ | 5421 | 5056 | 364.54 | 2.95 |
| TCC | _T_ | 7590 | 5500 | 2089.79 | 16.26 |
| AGG | _A_ | 7566 | 5263 | 2302.74 | 18.32 |

**Table 4:** Posterior coverage of the four parameters of the dynamic Ising model at four different T-mer sizes. Shown are the proportion of runs in which the true value fell within the 95% credible interval, for T-mers of length 4, 5, and 6 with overhangs of length 1, 1, and 2 respectively. The total number of runs was 61 for (6, 2, 2) T-mers and 121 for the others.

|  | (4,1,1) | (5,1,1) | (6,2,2) |
| --- | --- | --- | --- |
| $\lambda_-$ | 0.61 | 0.63 | 0.96 |
| $\lambda_+$ | 0.87 | 0.86 | 0.85 |
| $\beta$ | 0.76 | 0.81 | 0.94 |
| $\gamma$ | 0.69 | 0.76 | 0.85 |

We constructed models from three, nested, sets of mutation motifs. All models are strand-symmetric.

#### (basic)

Single-base changes, with separate rates for transitions (*μ_v_*), A↔T (*μ_AT_*), C↔G (*μ_CG_*), and other transversions (*μ_r_*):

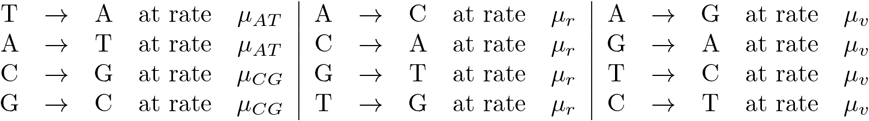

#### (CpG)

In addition to the above, an additional CpG rate:

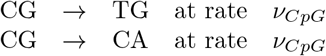

#### (repair)

In addition to the above, we included a collection of mutation motifs associated with known DNA lesion/repair pathways curated from the literature [reviewed in Sale et al., 2012, Goodman and Woodgate, 2013, Roberts and Gordenin, 2014, Cobey et al., 2015, Goodman, 2002]: deamination due to repair of UV-induced pyramidine dimers [*μ_UV_*, Sale et al., 2012, Sinha and Hader, 2002]; repair of guanine adducts at CpG sites Pfeifer [*μ_g_*, 2006]; deamination by AID [*μ*_AID_, Teng and Papavasiliou, 2007, Kasar et al., 2015]; deamination by members of the APOBEC family [*μ*_APOBEC_, Alexandrov et al., 2013a]; and repair by error-prone polymerases pol *η* [*μ_η_*, Alexandrov et al., 2013a]; and pol *ι* [*μ_ι_*, Roberts and Gordenin, 2014, Maul et al., 2016]. These are denoted here using ambiguity codes R: A or G, Y: C or T, W: A or T, and S: C or G; so for instance “TCW→TTW” means that both TCA→TTA and TCT→TTT share the same mutation rate.

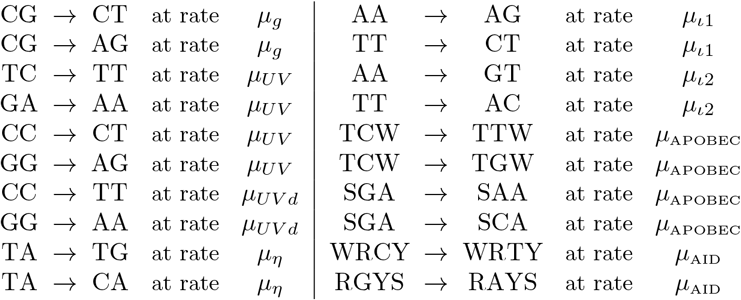

The mutational motifs included above are far from complete – context-specific affinities and error rates *in vivo* are not known for most DNA repair pathways – but neither do they exhaust the hypotheses available in the literature. For instance, many enzymes act more than once locally when they act, such as AID [Senavirathne et al., 2015] or error-prone polymerases involved in repair [Maul et al., 2016].

Nevertheless, we find significant effects for a number of biochemically-informed mutation features (Figure 5). Specifically, we find strong CpG and UV damage effects [Harris, 2015, also seen by]. It may seem tenuous to look for mutational signatures associated with cancer in the germline, or surprising that UV damage could play a significant role, but it is reasonable that shared repair machinery might induce similar context-dependent errors. Other likely sources of error in the germline include the effects of recombination [Arbeithuber et al., 2015, Myers et al., 2010].

**Figure 5:**
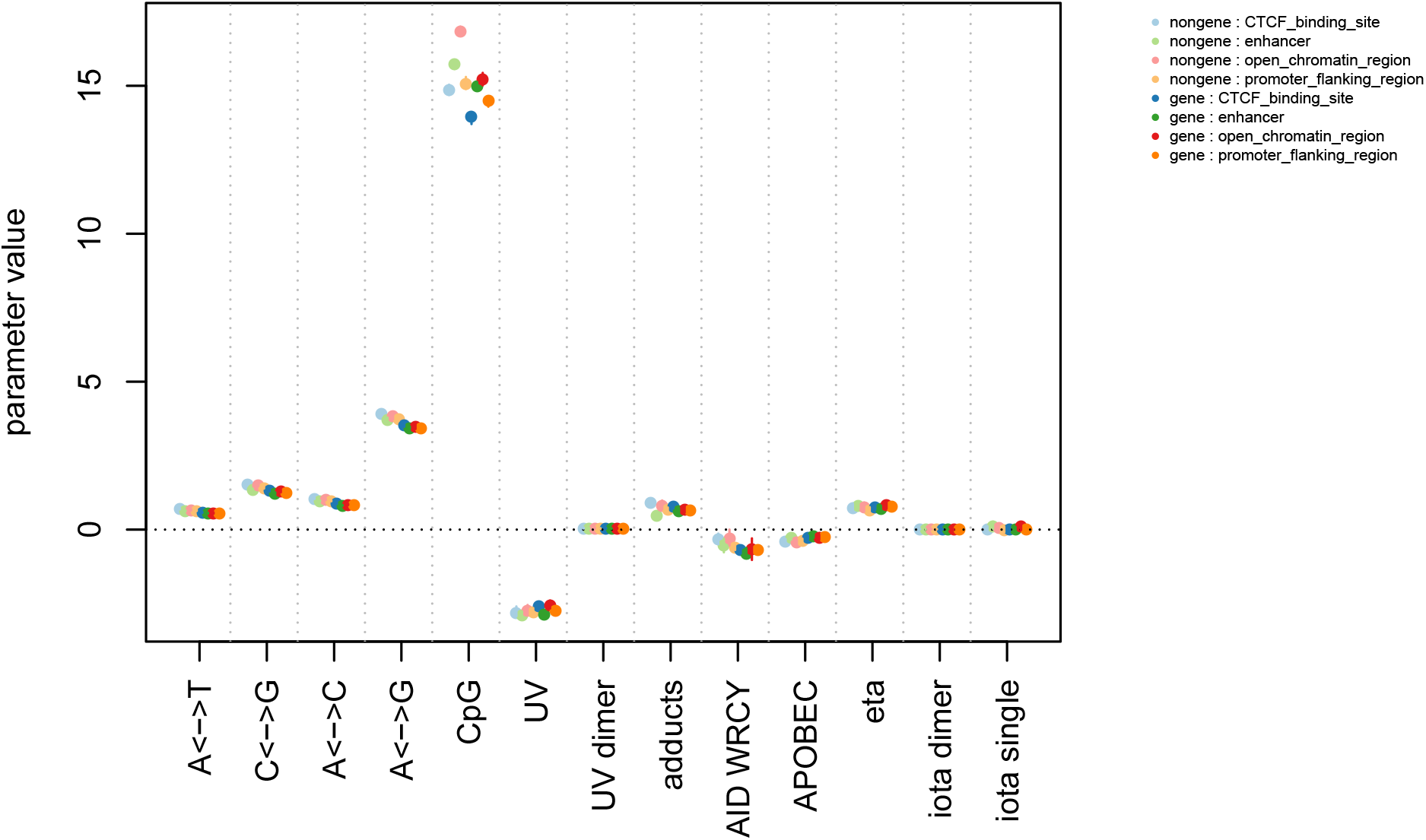
Estimated average mutation rates for the motifs in the “Hominid data” section since the common ancestor of humans and chimpanzee. Rates are scaled so that the mean time to common ancestor is 0.001 units – if we take this to be 6 million years [Scally et al., 2012, Langergraber et al., 2012] then rates are in units of mutations per 6,000 years. Shown are posterior medians (points) and 95% credible intervals (lines, which are quite small in all cases). Figure S2 shows the smaller rates in more detail, and Figure S3 shows the rates between categories relative to each other.

## 8 Discussion

This paper presents a tractable method for doing statistical inference using observations of a particle model in which probabilities of change are affected by nearby sites. Such models appear in many fields, for instance, as the dynamic counterparts to Markov random fields or spin glass models, and models of DNA sequence evolution.

The methods described here provide a way to efficiently approximate full likelihoods, which in many circumstances provide the optimal basis for inference [Neyman and Pearson, 1933], fast enough to allow Bayesian inference from Monte Carlo sampling. The method is fast thanks to its dependence on only local neighborhoods (so the time does not scale with sequence length once T-mer abundances are counted) and its reliance on sparse matrices whose structure can be precomputed and cached. It avoids approximations made by previous methods thanks to the use of T-mers and explicit quantification of the necessary length of the “arms” of the T-mers. We first discuss possible uses of the method, followed by challenges (and possible solutions).

### Other applications

One class of applications stems from *inference of process* – where the numerical values of various mutation rates are directly of interest. For instance, CpG hypermutability depends on the CGpair being methylated, an epigenetic modification that has important consequences for gene regulation; therefore, inference of CpG mutation rates in ancient taxa provides an indirect estimate of methylation status in extinct organisms. Similarly, many mutational motifs are thought to be caused by particular enzymes; inference of relative rates of these motifs may provide clues as to the activity and evolution of those enzymes. Furthermore, iterative model building by examining residual motif counts could provide a powerful way to discover unknown processes whose action had been obscured by other sources of noise.

In other applications, reconstruction of an unknown sequence may be of interest. For instance, training data might be used to estimate parameters of a noisy transmission line, facilitating the later reconstruction of noisily transmitted signals. Alternatively, multiple observations of independently transmitted copies of a single sequence (e.g., down a phylogenetic tree) might be used to estimate the process, and from this infer the original (ancestral) sequence (or sample from its posterior distribution).

### Limitations

The model presented assumes that the same process occurs at every location along the sequence, and that the length of the sequence does not change (no insertions or deletions). Neither of these assumptions are likely to be true for any data set of DNA sequence; however, both assumptions are also commonly made in phylogenetics. Some relaxation of the assumptions is possible – for instance, we can easily allow heterogeneity between sites by scaling rates by an overall gamma-distributed random factor [Yang, 1994]. Insertions and deletions are problematic, however.

Furthermore, we have not been able to put quantitative bounds on how well we can approximate the full likelihood with a given T-mer shape. One reason this is difficult is that there are special cases of initial and final sequence for which the likelihood is *not* approximable by looking only at T-mer counts; however, we believe such situations to be rare, so that the full likelihood can be well approximated on a set of outcomes of high probability. The TASEP model represents in some ways the worst-case of dependency propagation, and has an analytic solution [Schütz, 1997]. Further theoretical work in this direction would be welcome. In practice, the approximation may be checked by computing the likelihood using successively longer T-mers, and checking for convergence as the length (of both base and overhang) increase.

### Computation

The main factor that determines computation time is the lengths of the T-mers used for inference, since all computations scale with the number of possible patterns at the long end of the T-mer. Although the model is efficient enough to deal with fairly long T-mers, this exponential growth puts hard constraints on the application of the method. The length of T-mers required to accurately compute the likelihood in most models grows at most linearly with the mean density of substitution between the initial and final sequence. This does not in practice greatly increase the length of required T-mer since substitution densities approaching 100% are unlikely to be usable in practice especially if how the sequences align with each other is unknown. A more serious determining factor of T-mer length is the degree of dependency in the model itself. For instance, inserting the 13-base pair motif selected against by meiotic drive in primates [Myers et al., 2010] would require T-mers of width at least 25, and hence vectors of length 4^25^ ≈ 1.13 × 10^15^.

### Modeling

Careful building of complex models in this framework may take care, but no more than in any other class of models (e.g., multivariate linear regression). It is easy to specify nonidentifiable models – for instance, all 2mer mutation motifs include the 1mer motifs. The large number of possible mutation motifs also makes multiple testing an issue; an unguided approach might include all possible motifs but place a shrinkage prior on their values [Bhadra et al., 2015]. The converse approach of adding motifs that are overrepresented in the residuals may be quicker, but entails some degree of arbitrariness in choosing the motifs to add (and in which order). Finally, inference on trees may be difficult unless the root placement and distribution are strongly constrained: otherwise, initial investigations suggest the likelihood surface is often strongly ridged – for instance, increasing the abundance of G nucleotides at the root while also increasing the mutation rate away from G can achieve roughly the same result. However, in phylogenetic applications outgroups can be used to constrain these quantities.

In the genomic context, if we knew the specific mutational spectrum of each possible repair or copying pathways, then this framework could be used to rigorously estimate their average usages and impacts. Despite substantial progress over the past two decades [Goodman and Woodgate, 2013, Sale et al., 2012], this is still far from known. A promising alternative would be to use the distinct signatures obtained by dimension reduction techniques as in Alexandrov et al. [2013b], [Mathieson and Reich, 2017] or [Shiraishi et al., 2015] as a proxy for unknown but distinct pathways.

## 8.1 Acknowledgements

The initial impetus for this project came from Matt Dean; along the way substantial encouragement and useful suggestions were provided by Simon Tavaré, Rasmus Nielsen, Graham Coop, Yaniv Brandvain, and Sergey Nuzhdin. Many thanks are also due to Jessica Crisci for curating the human-chimp data. The majority of this work was done while PR was at USC.

FAM’s work supported by NIH grants R01 AI146028 and U01 AI150747. Dr. Matsen is an Investigator of the Howard Hughes Medical Institute. PLR’s work was supported by NIH award R01HG010774

## A Supplementary figures

**Figure S1:**
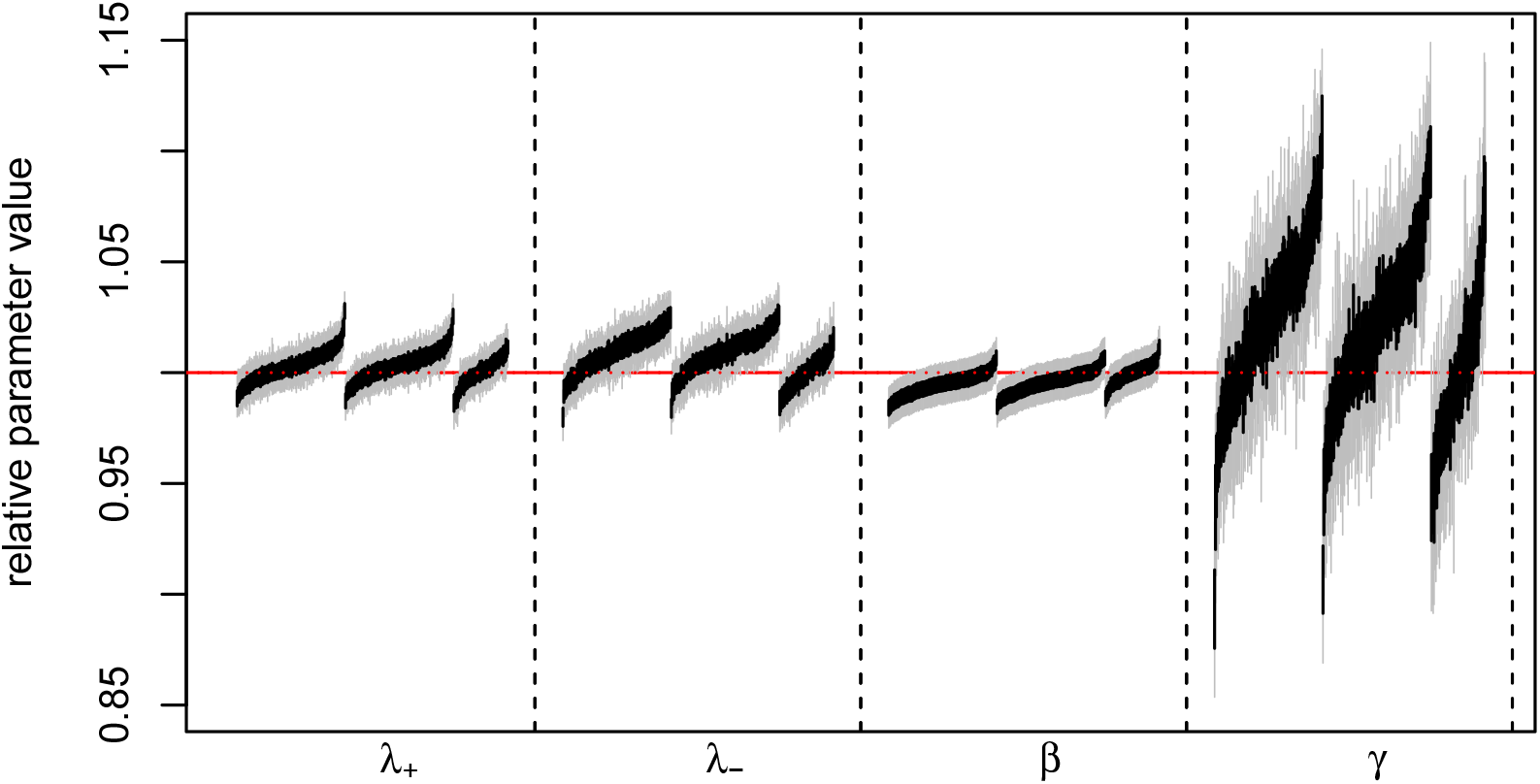
As in Figure 4: Credible intervals for the four parameters of the dynamic Ising model, from MCMC runs on independently simulated sequences of 10^6^ bases differing at around 18.6% of the sites. Shown are relative intervals, obtained by dividing by the true value. The three groups are left to right, credible intervals obtained from (4, 2,1), (5, 3,1), and (6, 2, 2) T-mers: note that only the longest (6, 2, 2) T-mers provide well-calibrated coverage (Table 4) and do not show bias.

**Figure S2:**
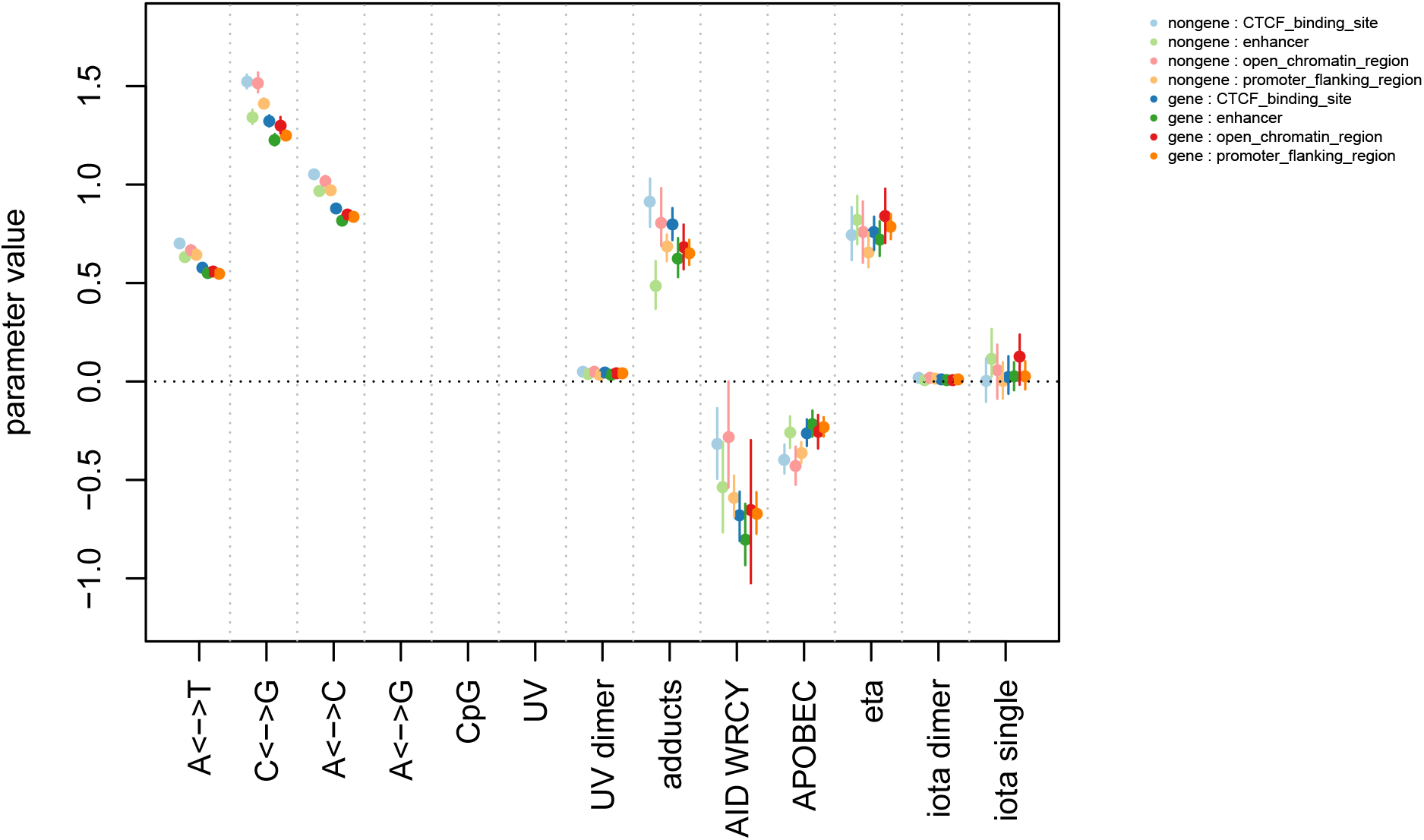
As in Figure 5, but with reduced vertical axis range to see in more detail: estimated average mutation rates for the motifs in the “Hominid data” section since the common ancestor of humans and chimpanzee. Rates are scaled so that the mean time to common ancestor is 0.001 units – if we take this to be 6 million years [Scally et al., 2012] then rates are in units of mutations per 6,000 years. Shown are posterior medians (points) and 95% credible intervals (lines).

**Figure S3:**
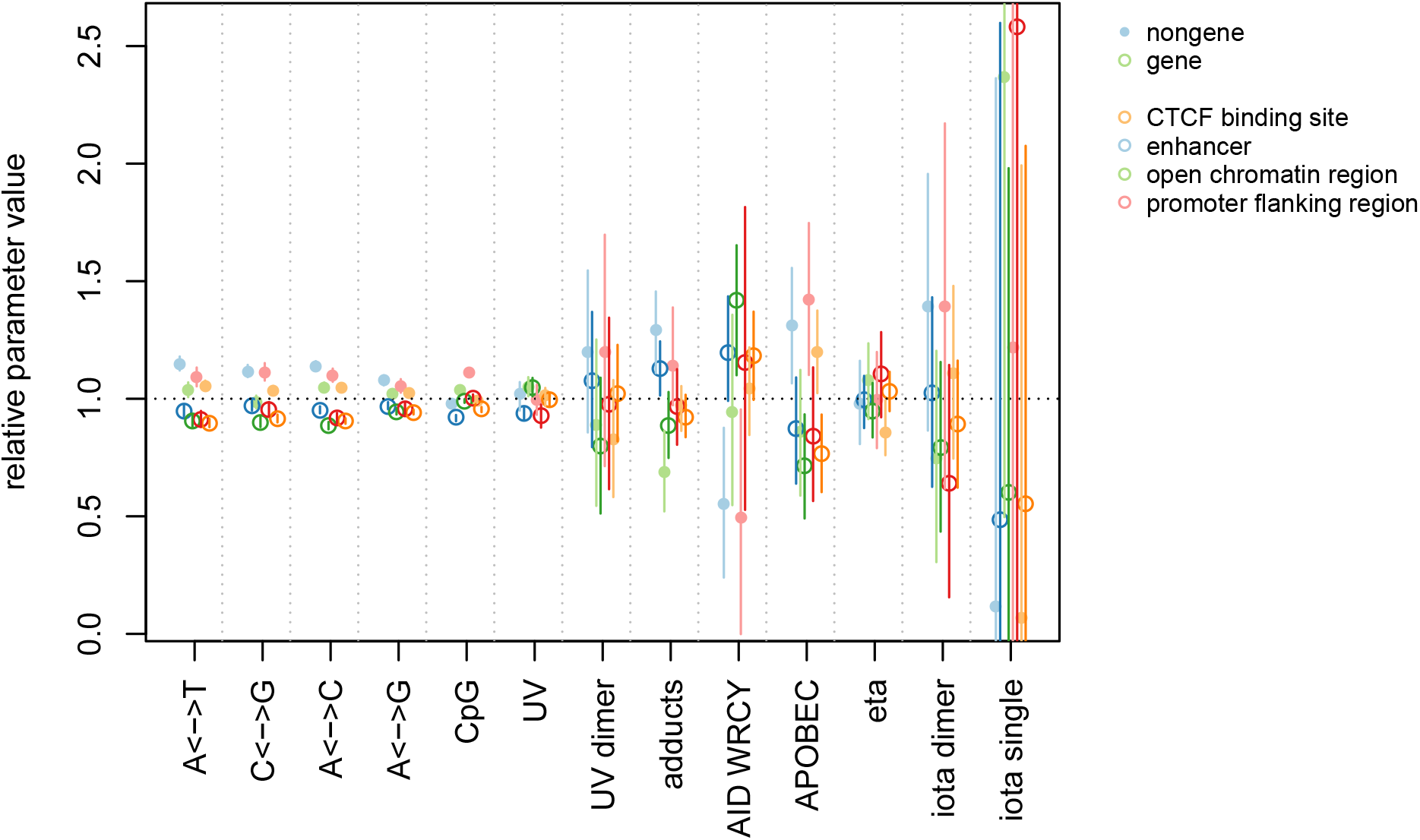
As in Figures 5 and S2, but where each rate is divided by the mean rate across all eight categories (as well as the 95% credible interval endpoints).

## B Proof of the approximation

First we quantify and prove the approximation (4). The basic idea is that the probability is only in error if there is a chain of mutations that create a dependency between the base of the T-mer and the flanking sequence. To simplify the argument, first assume that all transition triples change only a single, central site in the pattern, taking into account context on each side of maximum length *w*. (For instance, a trinucleotide model in which the flanking sites on each side affect the mutation rate of the central site has *w* = 1.)

Recall that, as described in the “Simulation” section, we may represent the process by first laying down a homogeneous Poisson process with rate *μ*\* per site of “potential mutations”, where *μ*\* is the maximum possible mutation rate, and then resolving these appropriately. (In particular, the times and locations of these potential mutations are chosen uniformly.) Suppose a possible change occurs at site *i* – *w*, at time *t*, which therefore might result in a change in mutation rate at site *i*. To resolve this, we need to know the outcome of any other potential changes that might change the sequence between *i* – *w* and *i* + *w*. Consider those that might happen only to the right of *i*, thus extending the sequence on which the change depends to the right. The number of such potential changes is Poisson with mean *μ*\* *wt*; if this number is nonzero, then this extends the amount of sequence on which the change at site *i* depends by a uniform number between 1 and *w*. If one occurs, say at time 0 < *s* < *t*, then any further potential changes that occur between time 0 and time *s* in the *w* sites to the right of this older potential change will again extend the range of dependency. Putting this together, the amount of sequence to the right of a site on which a mutation at that site depends is bounded by the compound Poisson process 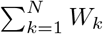, where *N* is Poisson with mean *μ*\**wt* and each *W_k_* is independent and uniform on {1, 2,…, *w*}.

Now consider the probability of interest,

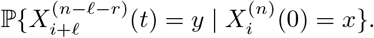

This probability can be written as a sum over all possible arrangements and resolutions of potential changes. Our approximation, *p_n,ℓ,r_*(*x,y*) is the probability that would be obtained by ignoring any potential changes that rely on context outside of the *n*-sequence *x*. So, if A is the event that the final subsequence *y* depends on sequence outside of *x* due to potential changes, then

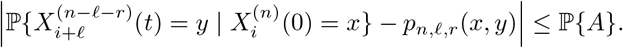

Since the size of the dependency window is a compound Poisson process, we can bound 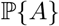 using the following lemma:

#### Lemma 1.

*Let* 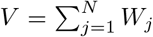, *where N is Poisson with mean* λ *and each W_j_ is independent and bounded between 0 and w. Then for k* ≥ 1,

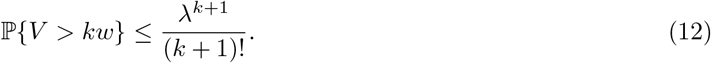

*Proof.* First note that 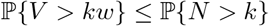. Now, use the fact that if *f*(*n*) = *n*(*n* – 1) ⋯ (*n* – *k*)/(*k* +1)! then *f*(*n*) = 0 for integers 0 ≤ *n* ≤ *k* and *f*(*n*) ≥ 1 for *n* > *k*, and so 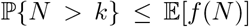. Now, 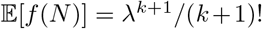 via direct calculation or the formula for the *k* + 1st factorial moment for the Poisson distribution.

Putting this together, we find that

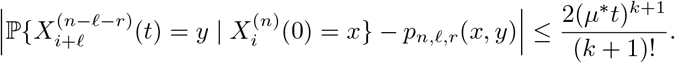

Here, the factor of 2 comes from the two sides (to the left and right) of the base of the T-mer.

The argument applies to more general transition triples, with *w* set equal to the maximum distance from a position that is changed to the edge of the context across all transition triples.

## C Pruning algorithm

Fix one tip of the tree, *a*, to be the taxa where we count “long” sequences, let *ρ* be the root, and let *b*_1_,…, *b_n_* be the remaining tips, where we observe “short” sequences. Let *v*_0_ = *ρ*, *v*_1_,…, *v_ℓ_* = *a* be the path from the root down to *a*, let *u*_1_, *u*_2_,…, *u_m_* be the remaining internal nodes, and for each *v_i_* let *u*(*v_i_*) be the offspring of *v_i_* that is not one of the other *v*. For each node in the tree, we will compute a matrix of probabilities: for nodes not on the path from a to the root, we compute the chance that that each “long” sequence observed at that node is matched by each possible combination of “short” sequences on the tips of the tree below it. For nodes on the path from a to the root, we compute the probability of observing each “long” sequence at that node, along with the combinations of “short” sequences on all tips further away from *a* (i.e., below that node if it was rooted at *a*). For a node *w* this will be stored as a 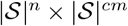 matrix, denoted *M*(*w*), where c is the number of tips for which *w* lies on the path from *a* to the tip. Rows of *M*(*u_j_*) sum to 1, while each entire matrix *M*(*v_i_*) sums to 1; the difference being that *M*(*v_i_*) sums over the prior at the root, and *M*(*u_j_*) takes the value at *u_j_* as given. Below, we index columns of *M*(*u*) by lists of short patterns, and leave the specific ordering to implementation. (For instance, if *c* = 2 then a generic entry is *M*(*u*)_*x*,(*y,z*_), where the long pattern *x* indexes the rows and the pair of short patterns (*y, z*) indexes the columns; in the software all matrices retain column names making this unambiguous.)

Let *t*(*w*) be the length of the branch above node *w*, for *w* ≠ *ρ*, and denote by *G^T^* the transpose of *G*.

1. Set *M*(*b_i_*) = *U* for each *b_i_*.
2. While there are *u_j_* whose children *w*_1_ and *w*_2_ both have matrices computed, let

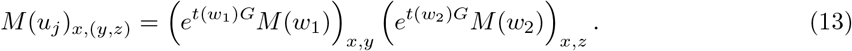
3. Let

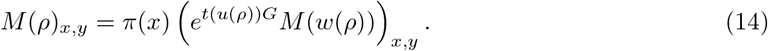
4. For each 1 ≤ *k* < ℓ, let

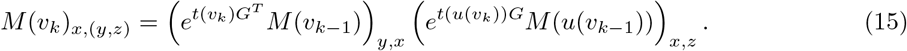
5. Finally, let

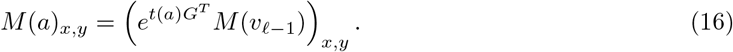

## Notes

https://github.com/petrelharp/context

### Competing Interest Statement

The authors have declared no competing interest.

